# Transformation from auditory to linguistic representations across auditory cortex is rapid and attention dependent for continuous speech

**DOI:** 10.1101/326785

**Authors:** Christian Brodbeck, L. Elliot Hong, Jonathan Z. Simon

**Affiliations:** Institute for Systems Research, University of Maryland, College Park, Maryland 20742, U.S.A; Department of Psychiatry, Maryland Psychiatric Research Center, University of Maryland School of Medicine, Baltimore, Maryland 21201, U.S.A; Department of Electrical and Computer Engineering, University of Maryland, College Park, Maryland 20742, U.S.A; Department of Biology, University of Maryland, College Park, Maryland 20742, U.S.A

**Keywords:** cohort model, phoneme surprisal, cohort entropy

## Abstract

During speech perception, a central task of the auditory cortex is to analyze complex acoustic patterns to allow detection of the words that encode a linguistic message. It is generally thought that this process includes at least one intermediate, phonetic, level of representations [1–6], localized bilaterally in the superior temporal lobe [7–10]. Phonetic representations reflect a transition from acoustic to linguistic information, classifying acoustic patterns into linguistically meaningful units, which can serve as input to mechanisms that access abstract word representations [11–13]. While recent research has identified neural signals arising from successful recognition of individual words in continuous speech [14–17], no explicit neurophysiological signal has been found demonstrating the transition from acoustic/phonetic to symbolic, lexical representations. Here we report a response reflecting the incremental integration of phonetic information for word identification, dominantly localized to the left temporal lobe. The short response latency, approximately 110 ms relative to phoneme onset, suggests that phonetic information is used for lexical processing as soon as it becomes available. Responses also tracked word boundaries, confirming previous reports of immediate lexical segmentation [18,19]. These new results were further investigated using a cocktail-party paradigm [20,21] in which participants listened to a mix of two talkers, attending to one and ignoring the other. Analysis indicates neural lexical processing of only the attended, but not the unattended, speech stream. Thus, while responses to acoustic features reflect attention through selective amplification of attended speech, responses consistent with a lexical processing model reveal categorically selective processing.

## Results and discussion

Magnetoencephalography (MEG) responses to continuous, narrative speech were analyzed with a framework designed to measure acoustic and lexical processing simultaneously. Source-localized brain responses were modeled as a linear filter response to multiple predictor variables that reflect acoustic and lexical properties of continuous speech (see Figure 1). Each source’s response time-course was modeled as a sum of responses to all predictors, such that all predictors competed for explaining variance in the response [15].

**Figure 1.**
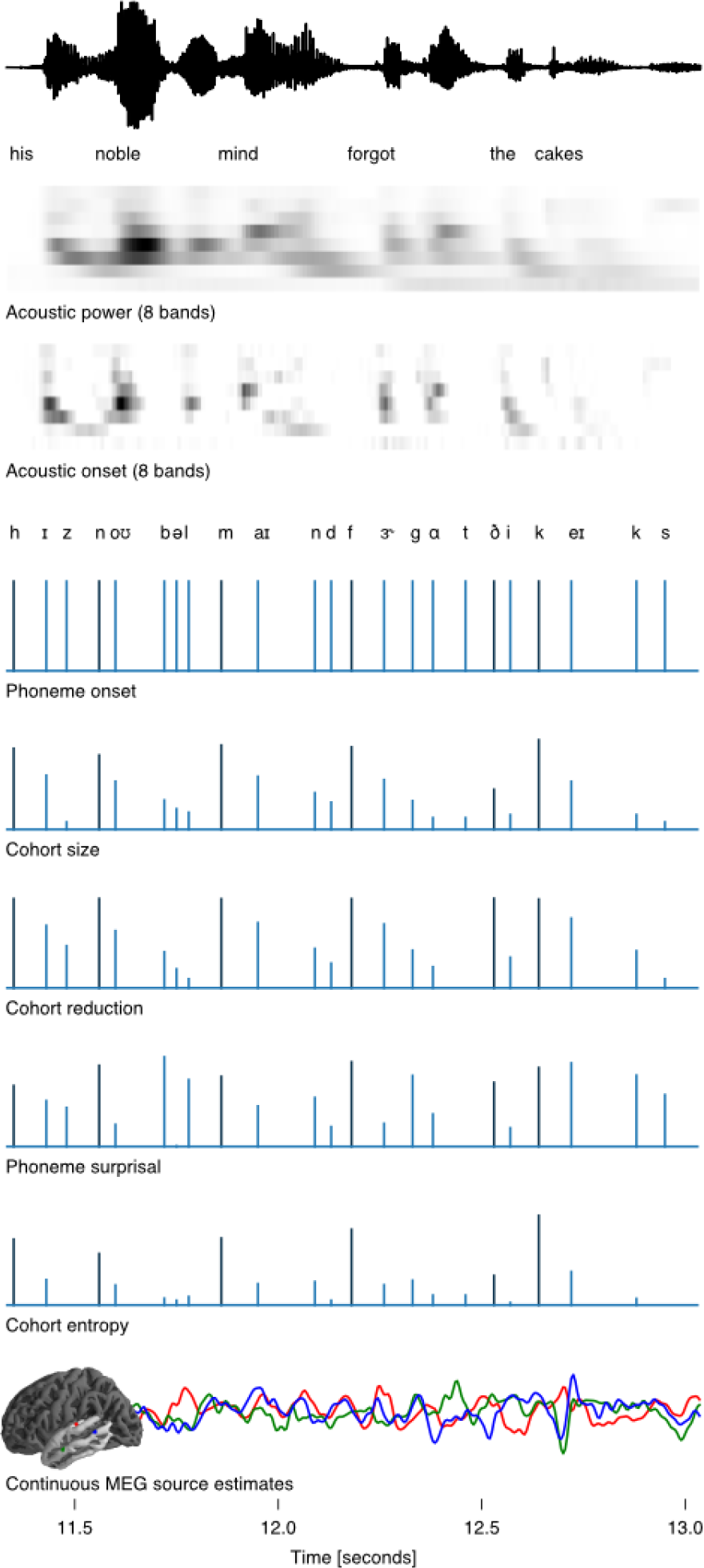
Analysis framework, illustrated with an excerpt from one of the stimuli. The acoustic waveform (top row) is shown for reference only. Subsequent rows show the predictor variables used to model responses to a single speaker. Acoustic predictors were based on an auditory spectrogram aggregated into 8 frequency bands. For the phoneme-based predictor variables, the initial phoneme of each word is drawn in black, whereas all subsequent phonemes are drawn in blue. The last row contains estimated brain responses from three virtual current dipoles, representative of the modeled signal. The anatomical plot of the cortex is shaded to indicate the temporal lobe, the anatomical region of interest (only the left hemisphere is shown, but both hemispheres were analyzed).

Acoustic properties were modeled through the envelopes of an 8 band auditory spectrogram, and their half wave rectified derivatives, to model both continuously varying **acoustic envelope energy** and **acoustic onsets** [22].

A binary **phoneme onset** variable was included to model responses to phonemes in general. Further variables with impulses of variable size at phoneme locations were used to model the modulation of phoneme responses due to lexical processing. All variables were based on the premise, widely supported by behavioral experiments, that phonetic information is used to incrementally constrain possibilities for the word that is currently being processed (e.g., [13,23,24]); this entails initial activation of multiple candidate lexical items, which are subsequently discarded when they become incompatible with the input. For example, after hearing the phoneme sequence /no⍰/, both *noble* and *notable* might be activated as potential candidates, but once the next phoneme /b/ becomes available (generating the prefix /no⍰b/), *notable* would be discarded as a possibility. This model suggests that at the occurrence of each phoneme in a word, there is a cohort of lexical items compatible with the current prefix. By using the word’s frequency in a large speech corpus [25] as proxy for its prior probability, a conditional probability distribution for the next phoneme can be computed at each stage in a word. The statistical variable **phoneme surprisal** associated with this probability distribution is a measure for how unexpected each phoneme is, based on the actual occurrence of the preceding phonemes, i.e., how much new, unexpected information it provides. **Cohort entropy** is defined as the Shannon entropy [26] over all lexical items compatible with the input at the given point in the word. Both phoneme surprisal and cohort entropy have previously been shown to be associated with reaction times [27,28] and MEG responses [29–32] to isolated spoken words. Related measures of **cohort size** and **cohort reduction** were also included in the framework because they are correlated with surprisal and entropy (see Table S1 and S2), but may be associated with different cognitive mechanisms. **Word onsets**, i.e. word-initial phonemes, were modeled separately from subsequent phonemes (black and blue in Figure 1), to account for the possibility that word onsets might involve different or additional processes, e.g., activation of an initial cohort as opposed to modification of an existing cohort (see also [32,33]).

### Responses to single speaker

MEG recordings from participants listening to a single talker were used to determine which variables were significant predictors of brain responses. Taken together, the 8 lexical processing variables significantly improved model predictions (*p* < .001). Because these variables are not independent (see Table S1 and S2), a first step consisted of reducing the initial set of variables to a set in which each variable explained a distinct proportion of the variance in the data. To this end, the significance of each lexical variable was evaluated, and the model was reduced by sequentially excluding non-significant predictors until only significant variables remained (cf. e.g. [27]). The model resulting from this procedure, henceforth called the baseline model, is shown in Figure 2; Figure 2 also shows results for non-lexical predictors in the baseline model. The left half of the figure shows anatomical plots, indicating where the predictor significantly improved predictions. Significance was assessed by comparing the predictive power of the baseline model to a model in which the predictor under investigation was shuffled in a way consistent with the appropriate null hypothesis. The right half of Figure 2 shows the filter kernels estimated for the baseline model, the so-called temporal response functions (TRFs). TRFs reflect the estimated response to an elementary temporal feature in the stimulus [34,35,15], and are thus a continuous analogue of evoked responses related to temporally distinct events. These responses were all estimated concurrently for the baseline model, i.e. they reflect each predictor’s contribution to the predictions of the model.

**Figure 2.**
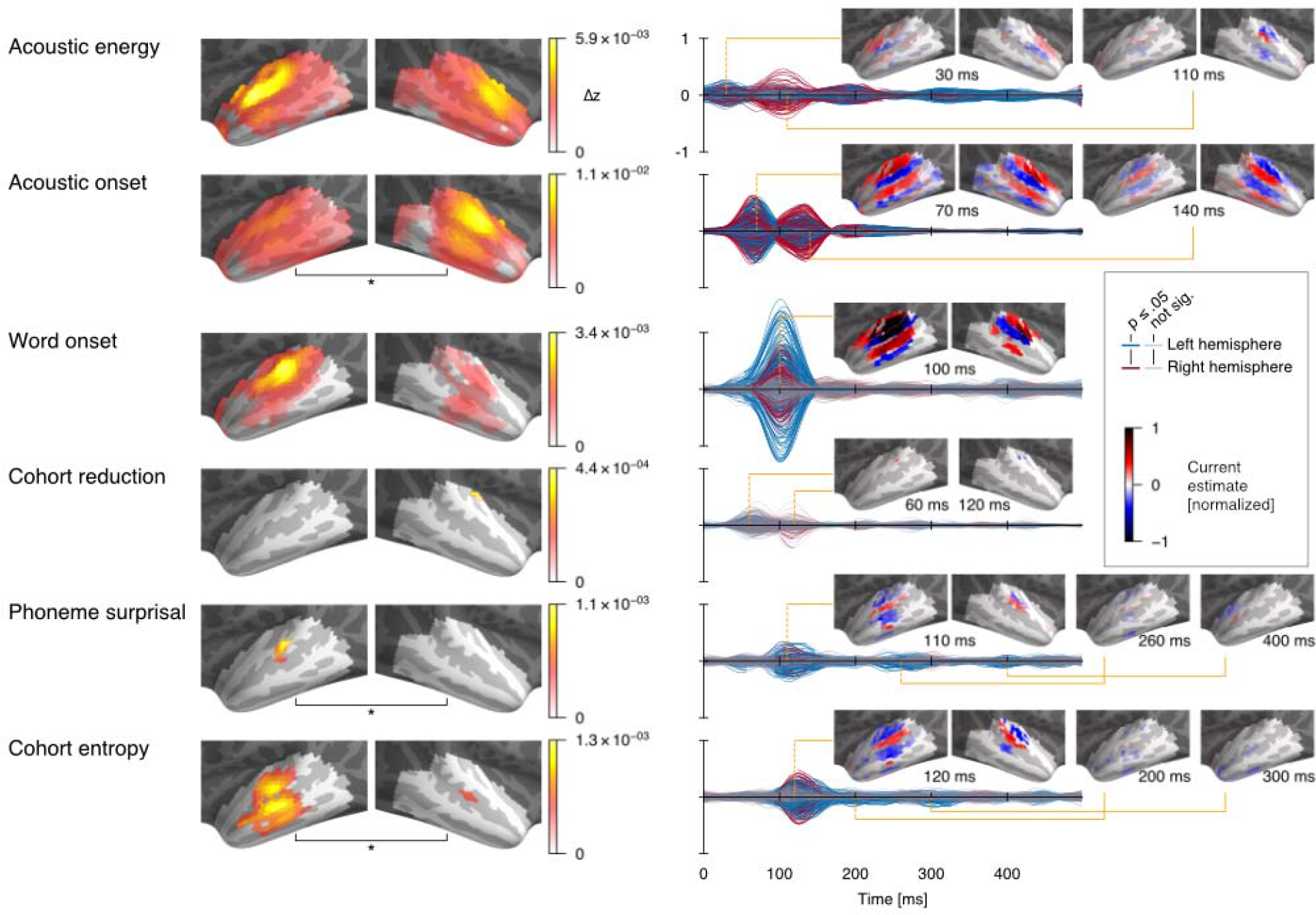
Brain responses to single speaker. Left column: significant predictive power (*p* ≤ .05 corrected). Colors reflect the difference in z-scored correlation between the full and the appropriately shuffled model. Color-maps are normalized for each predictor to maximize visibility of internal structure, as appropriate for evaluating source localization results: due to spatial dispersion of minimum norm source estimates, effect peaks are relatively accurate estimates, but strong effects can cause spurious spread whose amplitude decreases with distance from the peak. Right column: Temporal response functions (TRFs) estimated for the baseline model. Each line reflects the TRF at one virtual current dipole, with color coding its location by hemisphere, and saturation coding significance (*p* ≤ .05 corrected). Anatomical plots display TRFs at certain time points of interest (only significant values are shown), with color coding current direction relative to the cortical surface. Acoustic TRFs were averaged across frequency band for display as visual inspection revealed no major differences apart from amplitude differences between frequency bands. Effect peak locations are listed in Table S3.

The baseline model revealed significant left-lateralized contributions from both phoneme surprisal (*p* = .037, left > right *p* = .013) and cohort entropy (*p* < .001, left > right *p* = .028). The effect of surprisal was localized to core auditory cortex and nearby, whereas the effect of cohort entropy spread into more anterior and ventral regions, across the superior temporal sulcus, suggesting that the two variables reflect two different stages of speech processing. Consistent with this, the effect of surprisal peaked slightly earlier, at approximately 100-110 ms, whereas the effect of cohort entropy peaked at approximately 120-130 ms. This distinction is also consistent with the information that the two variables encode: Surprisal is a more local measure of phoneme prediction error [29], reflecting the frequency distribution of possible phoneme sequences up to the current phoneme. Cohort entropy on the other hand incorporates information about the frequency distribution over the cohort of lexical items that constitute possible continuations, possibly reflecting lexical competition [29,36]. More broadly, such activation of form and lexical item information in the superior and middle temporal lobe is consistent with reports of hemodynamic activity in this region [37–39], for example, effects of speech intelligibility [40] and generalization across different acoustic realizations of the same sentence [41]. Our results suggest that by 110 ms, *acoustic* information is used to update *phonetic* expectations held in the STG, and by 130 ms this update is used to constrain the activated cohort of *lexical* items. While the earliness of these effects might be surprising, evidence from gating studies suggests that 50-100 ms of input is sufficient to correctly identify the initial phoneme of a word [42], and it is plausible that the cortex uses this information as soon as it becomes available. Furthermore, these latencies are calculated from phoneme onset, without additional consideration of coarticulation cues. Since lexical processing is sensitive to coarticulation cues [5,43,44], information about phoneme identity may benefit from priming prior to the nominal phoneme onset.

None of the cohort-based predictors for word-initial phonemes remained in the model, suggesting that responses to initial phonemes could not be modeled by lexical distributions.

On the other hand, the effect of word-onset alone was highly significant (*p* < .001). Despite a numerically larger effect in the left hemisphere, lateralization was not significant (*p* = .087). Localization and TRF peak latency of this model variable were very similar to those of phoneme surprisal, suggesting that the two responses might be related to a shared mechanism. Indeed, word onset should be associated with disproportionately large surprisal, with little or no preceding phonetic information to constrain the cohort. A detailed examination of the TRFs suggests, however, that this explanation may not hold: the word onset TRF peak at 110 ms has the opposite polarity (i.e., current direction) from the corresponding peak in the surprisal TRF. One explanation consistent with this reversal in current direction arises from cohort theory, in which lexical candidates are activated at word onset, and then subsequently deactivated when a phoneme adds contradictory information [23]. In particular, when surprisal is high, candidates that were highly activated can be deactivated; when surprisal is low, expectations are confirmed and less change in activation levels is expected. A more general implication of this response is that word boundaries should be perceptually salient, despite the observation that clear cues for word boundaries are generally missing from speech waveforms [e.g. 45]. A similar word-onset electroencephalographic (EEG) response [19] emerged only after listeners learned to segment an artificial language into words [18], suggesting that it is not a response to local acoustic properties alone. A response tightly locked to word onset suggests that whichever cues listeners use to detect word boundaries [46–48], boundaries seem to be generally detected as they occur, rather than after incorporating cues occurring subsequent to word onset.

Both sets of acoustic predictors, envelopes and onsets, were associated with strong bilateral effects (both *p* < .001). Both were localized close to core auditory cortex, with acoustic onsets somewhat more predictive in the right hemisphere (lateralization *p* = .035). Significant effects extended over much of the temporal lobe, though the extended area could be due to spatial dispersion of MEG source estimates rather than genuine responses outside of core auditory regions [15]. Overall, the acoustic onset responses generate larger predictive power than the envelope responses, and thus may have absorbed some of the variance usually attributed to acoustic envelope representations when onsets are not considered [35,49]. TRFs to both sets of predictors exhibit two main peaks, consistent with earlier results [15,35,50]. Acoustic envelope energy was associated with peaks around 30 and 110 ms, of approximately opposite current direction. The latency of the pair of analogous peaks to the acoustic onsets is delayed, as expected due to the relationship between the two variables: The derivative operation shifts peaks in the acoustic energy towards earlier time points (the time of maximum rising slope precedes the time of maximum amplitude), thus increasing the distance between peaks in the predictor variable and specific time points in the neural response. The presence of analogous peaks in the TRFs to both acoustic representations might indicate that they jointly arise from a single, more complex underlying neural response type, reflecting both onset and continuous acoustic properties [51,52]. On the other hand, spatially, the effect of acoustic onsets localized posterior to acoustic envelope energy, which might instead indicate that the two responses stem from distinct neural populations that do not overlap completely.

The model also showed indications of a small effect of cohort reduction (*p* = .016), localized to the planum temporale, although the TRFs barely reached significance.

### Responses to two concurrent speakers

A reduced framework, consisting of the significant predictors for responses to a single speaker, was then used to model acoustic and lexical processing in a version of the cocktail-party paradigm [20,21]. Participants listened to a single-channel acoustic mixture of a male and a female speaker, attending to one and ignoring the other. This made it possible to specifically test whether the lexical processing observed for a single speaker is restricted to the attended speech stream, or whether it occurs for both attended and unattended streams. Figure 3 shows the predictive power of groups of predictors modeling relevant processing stages, and TRFs for the full model fitted to the two-speaker data.

**Figure 3.**
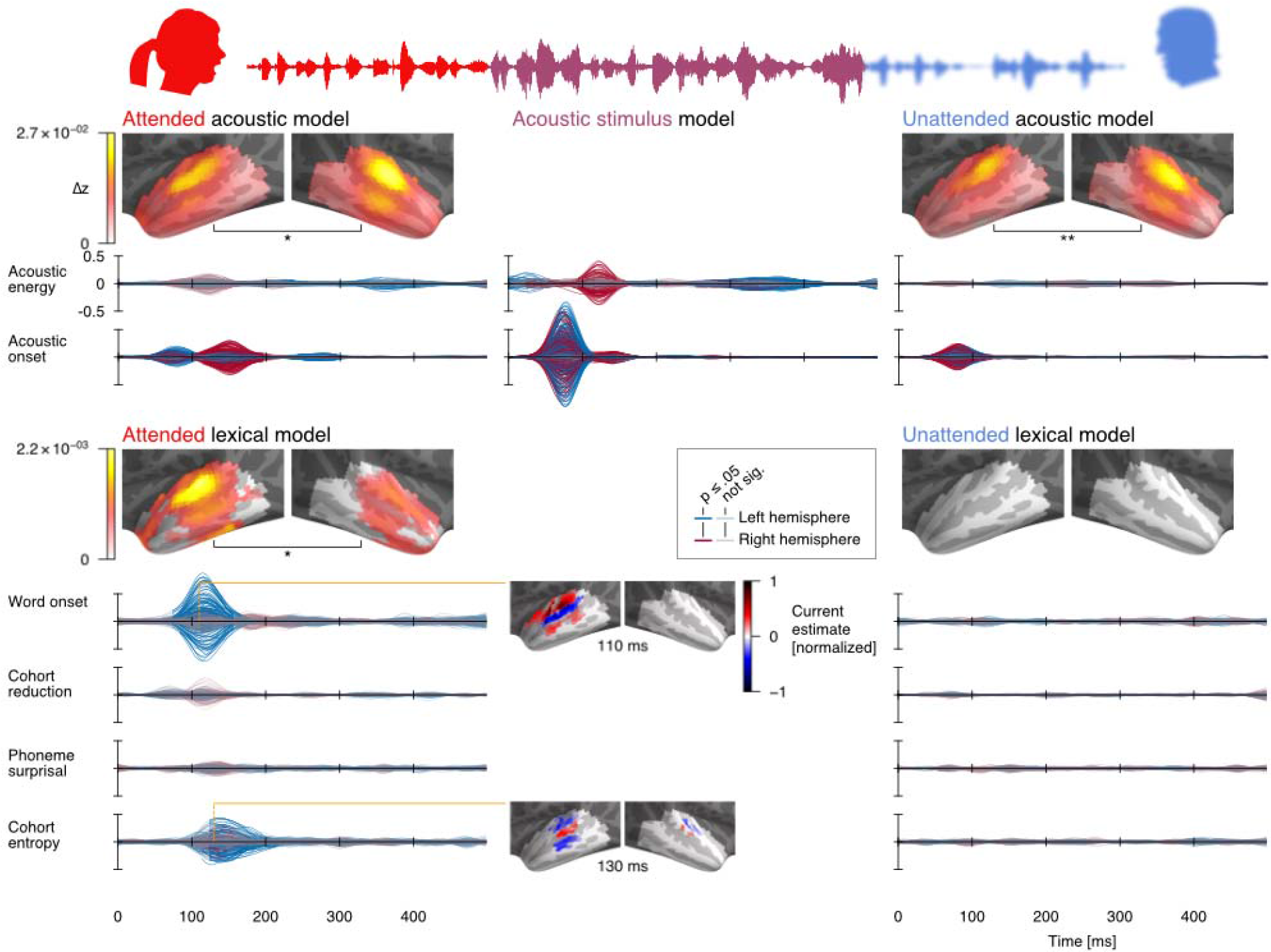
Brain responses to two concurrent speakers. Details analogous to Figure 2. The three columns display results for the model components for: the attended speech stream (left), the actual acoustic stimulus mixture (middle), and the unattended speech stream (right). The upper part of the figure displays results for acoustic features, the lower part for lexical processing.

Responses were significantly modulated by acoustic features of both the attended and the unattended speaker (both *p* < .001; lateralization *p* = .031 and .002). The relative amplitudes of the TRF peaks to acoustic onsets were consistent with previous results [35,50,53], with an earlier (~ 70 ms) peak predominantly reflecting the raw acoustic mixture (without selective enhancement of either speaker), and a later (~ 150 ms) peak predominantly reflecting acoustic energy in the attended speech. The TRFs to acoustic envelope energy almost exclusively reflected processing of the acoustic mixture, suggesting that auditory stream segregation may be predominantly reflected in onset processing.

In contrast to the acoustic models, only the lexical processing model for the attended speech showed significant effects (*p* < .001); lexical properties of the unattended stream did not (*p* = .275), and the effect of lexical processing of the attended speech was significantly greater than for unattended speech (*p* < .001). TRFs also supported this conclusion, with significant response to word onset and cohort entropy in the attended speech only. These responses were very similar to the corresponding single speaker responses, although with reduced amplitude, and an additional delay of ~ 10 ms.

This finding suggests that well-known interference effects from meaningful content of unattended speech [20,54–56] do not arise from lexical processing of the unattended stream. While recent research suggests that processing of information contingent on successful word recognition is suppressed for unattended speech [14,16], these findings left open the possibility that unattended speech is processed up to and including identification of lexical items, but without retrieval of the recognized words’ properties. The results of the present investigation indicate that lexical processing of unattended speech is suppressed at the level of detecting word forms.

The absence of lexical responses to unattended speech raises the possibility that lexical processing constitutes a bottleneck in speech perception. Lexical perception is thought to be massively parallel by involving activation of multiple candidate lexical representations through the cohort [11]. The mechanisms implementing this multiple activation might involve mental resources that cannot be shared across multiple parallel instances of the same process, making it impossible for more than one cohort to be represented at the same time.

In sum, MEG responses to continuous speech reflect a transformation of the speech signal from acoustic representations, which can be characterized with spectro-temporal receptive fields, to probabilistically driven activation of lexical units. Phonetic cues are rapidly analyzed for their relevance to word perception, updating a lexical processor in the left temporal lobe within ~ 130 ms. In the presence of two competing speakers, this transformation is restricted to the attended speech stream. While the analysis presented here is naturally limited to a specific kind of listening condition, and adults with normal hearing, the framework introduced here lends itself to studying the influence of different conditions and individual differences on speech processing.

## Acknowledgements

This work was supported by a National Institutes of Health grant R01-DC-014085 (to JZS) and by a University of Maryland Seed Grant (to LEH and JZS). We would like to thank Krishna Puvvada for his assistance in designing and preparing the stimuli, and Natalia Malley for her help in collecting data and for excellent technical support.

## Author Contributions

J.Z.S. and L.E.H. conceived and performed the experiments and secured funding. C.B. and J.Z.S. conceived and performed the analysis and wrote the manuscript.

## Methods

### Participants

MEG data were recorded from 28 native speakers of English, recruited by media advertisements from the Baltimore area as control group for another study. Participants with medical, psychiatric or neurological illnesses, head injury, and substance dependence or abuse were excluded. All subjects signed informed consents and were paid for their participation. Data from two participants were excluded, one due to corrupted localizer measurements, and one due to excessive magnetic artifacts associated with dental work. The sample analyzed was composed of 18 male and 8 female participants with mean age 45.2 (range 22 - 61). All subject provided informed consent in accordance with the University of Maryland Baltimore Internal Review Board.

Three participants were left handers. Excluding them from analysis did not change the major results, but some individual effects changed significance. For responses to a single speaker, the effect of cohort reduction lost significance (*p* = .091), and, after dropping this variable from the model, the effect of bare phoneme onset became significant (*p* < .001, right > left *p* = .018). In the two-speaker model, now fitted without cohort reduction, the lateralization of the acoustic and lexical model for the attended speaker lost significance (*p* = .082 and *p* = .106).

## Method details

### Stimuli

One minute long segments were extracted from audiobook recordings of A *Child’s History of England by Charles Dickens*, one chapter read by a male and one by a female speaker (https://librivox.org/a-childs-history-of-england-by-charles-dickens/, chapters 3 and 8). Pauses longer than 300 ms were shortened to an interval randomly chosen between 250 and 300 ms, and loudness was matched perceptually. Cocktail party stimuli were generated by additively combining two segments, one form each speaker.

Four segments were extracted for each speaker: male-1 through 4 and female-1 through 4; mix-1 through 4 were constructed by mixing male-1 and female-1, and so forth. Participants listened four times to mix-1, while attending to one speaker and ignoring the other (which speaker they attended to was counterbalanced across subject), then 4 times to mix-2 while attending to the other speaker. Then, the four segments just heard were all presented once individually. The same procedure was repeated for stimulus segments 3 and 4. After each mix segment, participants answered a question relating to the content of the attended stimulus.

Participants lay supine and were instructed to keep their eyes closed during stimulus presentation (to minimize ocular artifacts and head movement). Stimuli were delivered through foam pad earphones inserted into the ear canal.

### MEG data acquisition and preprocessing

Continuous MEG data were acquired with the 157 axial gradiometer whole head MEG system (KIT, Kanazawa, Japan) inside a magnetically-shielded room (Vacuumschmelze GmbH & Co. KG, Hanau, Germany) at the University of Maryland, College Park. Sensors (15.5 mm diameter) are uniformly distributed inside a liquid-He Dewar, spaced ~ 25 mm apart, and configured as first-order axial gradiometers with 50 mm separation and sensitivity > 5 fT·Hz^−1/2^ in the white noise region (> 1 KHz). Data were recorded with an online 200 Hz low-pass filter and a 60 Hz notch filter at a sampling rate of 1 kHz.

Recordings were pre-processed using mne-python [57,58]. Flat channel responses were automatically detected and excluded. Extraneous artifacts were removed with temporal signal space separation [59]. Data were filtered between 1 and 40 Hz with a zero-phase FIR filter (mne-python 0.15 default settings). Responses time-locked to the onset of the speech stimuli were extracted and downsampled to 100 Hz.

### Source localization

Before the MEG recording, each participant’s head shape was digitized (Polhemus 3SPACE FASTRAK) and five marker coils were attached to their head. The marker coils were localized with respect to the MEG sensors at the beginning and at the end of the recording session, and these position measurements were used to determine the head position relative to the MEG sensors. The digitized head shape was used to coregister the FreeSurfer [60] “fsaverage” template brain to each subject’s head shape using rotation, translation and uniform scaling.

A source space was defined using four-fold icosahedral subdivision on the white matter surface of the fsaverage brain, with all source dipoles oriented perpendicularly to the cortical surface. Based on this source space, *ℓ*^2^ minimum norm current estimates [61,62] were computed for all data using depth weighting parameter of 0.8 [63]. Analysis was restricted to the temporal lobe of both hemispheres, based on anatomical labels in the “aparc” parcellation [64].

## Quantification and statistical analysis

### Predictor variables

Predictor variables were generated as uniform time series with a sampling rate of 100 Hz to match the processed MEG data. Figure 1 illustrates the predictor variables, aligned with an excerpt from one of the stimuli.

Responses to the acoustic features of the speech signal were modeled using a model of acoustic transformations in the brain stem, the so called auditory spectrogram [65]. The auditory spectrogram was computed using the NSL Toolbox (http://www.isr.umd.edu/Labs/NSL/Software.htm) and shifted by −20 ms in order to compensate for the intrinsic delay introduced by this transformation. A predictor reflecting moment by moment **acoustic envelope power** was generated by summing the auditory spectrogram in 8 logarithmically spaced frequency bands. Because brain responses are known to be sensitive to contrast and changes, and phonetic information is often specifically located in acoustic onsets [22], it was important to control for responses to such acoustic onsets in the acoustic signals. For this reason, an **acoustic onset** predictor was constructed from the half-wave rectified derivative of the acoustic envelope predictor.

All phoneme-based predictors were modeled as impulses at phoneme onset (see Figure 1). Phoneme onsets in the stimuli were automatically determined by the Gentle forced aligner (https://lowerquality.com/gentle/) and then adjusted by hand where appropriate. A phonetic lexicon with lexical statistics was generated by combining pronunciations from the CMU Pro-nouncing Dictionary (http://www.speech.cs.cmu.edu/cgi-bin/cmudict) and word frequency statistics from the SUBTLEX subtitle database [25]. Stress information was stripped from all phonemes. Missing pronunciations were manually added, and words occurring in the stimuli but missing from SUBTLEX were assigned a frequency count of 1.

The cohort refers to the set of words compatible with the acoustic input at any point during a word [11]. For each phoneme, the cohort was determined by selecting from the phonetic lexicon those entries that started with the phoneme sequence from the beginning of the word to the current phoneme. The **cohort size** variable was the log of the size of the cohort at each phoneme. The **cohort reduction** variable was the log of the number of words at the current phoneme minus the number of words at the previous phoneme or, for the initial phoneme, minus the number of words in the whole lexicon. While these two variables are not as widely used as surprisal and entropy (see below), they are potentially more fundamental variables that should be controlled for before drawing conclusions about surprisal and entropy.

While cohort size variables depend only on number of words, the frequency of individual words is known to affect lexical processing [13,66]. This is taken into account by the measures of phoneme surprisal and entropy. **Phoneme surprisal** is defined as the inverse of the conditional probability of each phoneme, given the preceding phonemes in the current word:

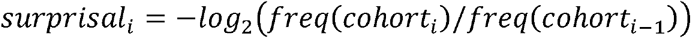

Where *cohort_i_* is the cohort at phoneme with position *i*, and *freq(c)* is the summed frequency of all words in cohort *c*. **Cohort entropy** is defined as the entropy [26] of the cohort at each phoneme. Entropy at phoneme *i* is given by:

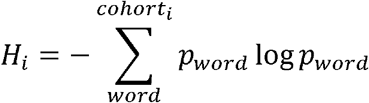

Where *p_word_* is the probability of the given word form, here modeled as the relative frequency.

To account for the possibility that the initial phoneme of each word is processed differently from the subsequent phonemes (see e.g. [11]), we modeled the initial phoneme of each word separately from the subsequent phonemes for each variable (indicated by a different color of the word-initial phonemes in Figure 1).

### Model estimation

For each subject, the localized current at each potential neural source dipole was modeled as a sum of linear convolutions of the stimulus variables with a filter of 500 ms duration. Optimal filters were estimated for all predictor variables concurrently with a coordinate descent algorithm [15,67] minimizing the *ℓ*1 error between predicted and actual current time course. Filters were generated from a basis of 50 ms Hamming windows, centered at each time point in the kernel. This smoothness constraint on the filters was imposed to improve the reliability of predictions, compensating for the temporal sparseness of the impulse representation of phonemes. Algorithms used for model estimation and statistical analysis are publicly available in the Eelbrain open source Python package [68] (https://github.com/christianbrodbeck/Eelbrain).

### Statistical analysis

Model fit was estimated using the *z*-transformed Pearson correlation between estimated and measured responses. Model fit *z*-maps were smoothed with a Gaussian kernel (STD = 5 mm) to account for granularity caused by local variation in source dipole orientation. To compare the fit of two models, a *t*-map was computed by applying a related measures *t*-test at each source dipole. The resulting map was processed with threshold-free cluster enhancement (TFCE) [69], and a distribution of the largest expected TFCE value per *t*-map under the null-hypothesis was computed with 10,000 permutations, randomly switching condition labels within subject [69,70]. A *p*-value for each dipole was computed by locating the original TFCE-value on the permutation distribution.

To test for significant contributions of a given predictor, the predictive power of the full model was compared to an alternative model, which was identical except for the predictor under investigation, which was shuffled in a way appropriate for the hypothesis being tested. A predictor was considered significant if it significantly improved model fit across participants. This procedure allowed testing for incremental model improvement due to a specific predictor, without introducing bias by changing the degrees of freedom. Under the null hypothesis that there is no significant association between the given predictor and the responses, a shuffled version of the predictor should be equally effective as the properly aligned version. A difference in model fit between the full and the shuffled model thus indicates a significant temporally specific relationship between predictor and responses. Table S3 lists for each significant effect the anatomical location of maximum model fit improvement.

Tests of hemispheric asymmetry were performed by comparing the model fit improvement between the two hemispheres. First, a difference map was computed by subtracting from the *z*-values of the full model those from the shuffled model. The resulting difference maps from both hemispheres were mapped to the left hemisphere of the “fsaverage_sym” brain [15,71], masked by the region of significant model improvement in at least one hemisphere, and compared with a two-tailed *t*-test while controlling for multiple comparisons with TFCE as described above.

Temporal response functions, i.e. the kernels of the optimal filters determined through boosting, were analyzed similarly, but including the additional dimension of time. A spatio-temporal *t*-map was computed for a one-sample *t*-test against 0. This map was again processed with TFCE and a two-tailed distribution for the maximum TFCE value was computed based on 10.000 permutations. For graphical display only, time series were upsampled to 500 Hz to minimize visual discretization artifacts.

### Single speaker analysis

Responses to a single speaker were used to determine variables that reflect lexical processing of phonetic information. To test for an effect of lexical variables without inflated type I error due to multiple comparisons, an initial test was performed against a shuffled model in which all 8 lexical variables were shuffled together. Subsequently, the set of lexical variables was reduced to a set in which each variable explained a distinct proportion of the variance. To this end, the model was reduced one predictor at a time by removing the predictor whose model contribution was least significant, until only significant predictors were left (see e.g. reference [27] for a similar approach). Once only significant lexical predictors remained (henceforth called the reduced model), the other variables in the model were also evaluated for significance (shown in Figure 2).

The way in which variables were shuffled depended on the nature of the variable and the corresponding null hypothesis: Lexical variables were shuffled by randomly reordering the values (e.g. phoneme surprisal) while leaving the phoneme time locations constant. Acoustic predictors were shuffled by swapping the first half (0-30 s) of each stimulus with the second half (30-60 s). To test the effect of word onset, word onsets were randomly assigned to phoneme locations, while keeping all phoneme locations constant. To test for the effect of phoneme location above word onsets, the time-series representing subsequent phonemes (excluding word onset phonemes) was swapped in the same manner as the acoustic predictors. In each case, all remaining predictors were left unchanged in the control model.

### Two speaker analysis

For modeling responses to stimuli with a mixture of two speakers, separate predictors were included for the attended and the unattended speech stream, both based on the reduced single speaker model. In addition, acoustic predictors were generated for the acoustic mixture of the two speakers, i.e., for the raw acoustic stimulus that was actually presented to the participants.

Models were assessed by grouping predictors modeling each process of interest. Since the acoustic mixture is closely approximated by a linear combination of attended and unattended signal (in all bands *r* > .95 for acoustic envelope energy and *r* > .88 for acoustic onsets), predictive power of the mix could not be assessed independently. Instead, predictive power of the attended stimulus was assessed by shuffling both attended and mix acoustic predictors, and the unattended stimulus was assessed by shuffling both unattended and mix acoustic predictors. Nevertheless, acoustic TRFs could be analyzed for all three streams, since the coordinate descent algorithm determines which predictor can reduce the error most efficiently, regardless of whether the same model fit could be achieved by a linear combination of other, less efficient predictors. Lexical processing was assessed separately for attended and unattended streams, by shuffling all the values among phoneme locations, but leaving phoneme locations themselves unchanged. Thus, the model comparison controlled for responses associated with all phonemes independent of lexical processing.

